# Haplotype-resolved genomes and population genetics to analyze glandular secretory trichome formation mechanism in oregano

**DOI:** 10.1101/2024.04.30.591903

**Authors:** Meiyu Sun, Jiahui Miao, Ningning Liu, Yanan Zhang, Jinzheng Zhang, Di Wang, Fei Xia, Hongtong Bai, Hui Li, Lei Shi

**Affiliations:** State Key Laboratory of Plant Diversity and Specialty Crops, Institute of Botany, Chinese Academy of Sciences, Beijing 100093, China; China National Botanical Garden, Beijing 100093, China; University of Chinese Academy of Sciences, Beijing 100049, China

**Author notes:** Correspondence: Lei Shi,; Hui Li. These authors contributed equally to this work (Meiyu Sun, Jiahui Miao).

## Abstract

Oregano is an important economic plant which has valuable medicinal and aromatic properties. Oregano essential oil, containing carvacrol and thymol, is a preferred material to replace antibiotics in feed additives. Glandular secretory trichome (GST) density has positively correlated with the production of these compounds. Here, two haplotype-resolved genomes were assembled and annotated which contained 15 chromosomes with the total length of 606.75 and 612.74 Mb, respectively. Oregano had experienced two whole-genome duplications corresponding to the divergence ∼5.120/4.564 and ∼66.857/69.923 Mya, respectively. Many transcription factors and genes were found related to GSTs formation mechanism such as R2R3-MYB- and HD-ZIP IV-encoding genes. 2,669,410 SNPs, 569,093 InDels, 14,839 DUPs, 110 INVs, 3,976 TRANSs, and 1,426 CNVs were detected among two haplotype-resolved genomes. Two high density genetic linkage maps consisted of 15 LGs and spanned 2,279.28 and 2,322.83 cM, respectively. GADS, GABS, and GTS of F_2_ segregating populations showed obvious superparental dominance. One/one, one/one, and two/four QTLs for GADS, GABS, and GTS were independently mapped on two genetic maps, respectively. Five candidate genes showed extreme difference in two bulked segregant pools. Our study not only provides significant insight into the GSTs formation mechnism, but also will facilitate molecular breeding in oregano.

**One Sentence Summary:** Oregano essential oil is a preferred material to replace antibiotics which will use to study the glandular secretory trichome formation mechanism and analyze the source of essential oil.

## Introduction

The genus *Origanum*, belonging to the Lamiaceae family, is widely distributed worldwide which contains more than 44 species and 7 subspecies of herbaceous perennials or sub-shrubs with valuable ornamental, flavoring, medicinal, feed additive, and aromatic properties (http://www.theplantlist.org/tpl1.1/search?q=Origanum). There is only one species *O. vulgare* L. which was recorded by Flora of China (Li et al. 1994). Among those oregano varieties, *O. vulgare* ‘Hot & Spicy’ is of particular interest for its high oil yield (2.80%) and high carvacrol content (89.90%). Since ancient times, oregano essential oils (EOs) have been extensively utilized in folk medicine due to their strong antibacterial properties against a variety of diseases (Marinelli et al. 2018; Hao et al. 2022). EOs’ antimicrobial and antioxidant therapeutic properties are dependent on their bioactive ingredients, which include thymol and carvacrol (Falleh et al. 2020; Lu et al. 2021). In fact, a number of plants in the Lamiaceae family including peppermint and thyme have found widespread use in the food industry. These plants are also used as food additives to prevent deterioration (Kang et al. 2019; Zhong et al. 2021). Oregano essential oil, containing carvacrol and thymol, is a preferred material to replace antibiotics and has become a research hotspot in feed additives (Wang et al. 2023). We hope to greatly improve the yield and quality of oregano EOs through molecular breeding.

Trichomes are widely distributed in different organs of plants including flowers, leaves, and stems, and are classified as either glandular or non-glandular trichome (Werker et al. 2000). Mint, basil, lavender, oregano, and thyme (Lamiaceae) are cultivated for the terpenoids produced in their glandular secretory trichome (Maleci et al. 2006). Glandular secretory trichome (GST) density has been proven to be positively correlated with the production of these compounds (Yan et al. 2017). Thus, boosting the density of GSTs might be a useful strategy to raise these natural metabolites’ productivity (Tissier et al. 2012). Many GSTs formation mechanism are mainly come from the works on *Artemisia*, tobacco, and tomato (Robert and Tissier 2020) including R2R3-MYB (Lv et al. 2022) and HD-ZIP Ⅳ (Xie et al. 2021) transcription factors, *HOMEODOMAIN PROTEIN 1* (*AaHD1*), *GLANDULAR TRICHOMESPECIFIC WRKY 2* (*AaGSW2*), *JA ZIM-domain 8* (*AaJAZ8*), *WRKY* transcription factor (*AaGSW2*) (Xie et al. 2021), as well as receptors involved in phytohormone-induced signaling cascades (Chen et al. 2023), etc. Therefore, the genetic network controlling GSTs formation is not only the main target to study the synthesis and regulation of secondary metabolites in plants, but also the key element to understand plant breeding strategy (Huchelmann et al. 2017; Chalvin et al. 2020). Genetic linkage maps have been employed in marker-assisted quantitative trait locus (QTL) investigations to analyze the genetic basis of complex traits in a variety of fruit and horticultural plants, including grape (Zhang et al. 2023), kiwifruit (Wang et al. 2023), crape myrtle (Zhou et al. 2023), etc. Single nucleotide polymorphism (SNP) markers are an effective method for creating high-density genetic maps for QTL analysis. As a result, they might be used in plants to more thoroughly investigate the genetic process underlying significant quantitative features. A few studies have identified QTL linked with GST density in plants having GST. Using an F_4_ mapping population and a genetic linkage map made from genotyping by sequencing (GBS) data and consisting of 2121 SNP markers, two main QTL regulating capitate glandular trichome (CGT) density in sunflower florets were identified (Gao et al. 2018). One significant QTL, tric11, controlled the density of glandular trichomes type I in melon and accounted for between 23.8 and 58.7% of its phenotypic variance (Palomares-Rius et al. 2016). Six QTLs on chromosomes I, VII, VII, and XI were identified by QTL mapping of the morphology of type VI glandular trichomes in tomato, and a 500 gene interval in an advanced population generated from one of the back-cross lines was found (Bennewitz et al. 2018). These studies pointed out the direction for the construction of oregano genetic population and high-density genetic linkage map, QTL mapping of GSTs, and the screening of key genes regulated to GSTs.

Hybridization between or within species can produce heterozygous genomes, which can contribute to the preservation of plant variety and perhaps lead to the emergence of new species (Baek et al. 2018). Precise characterization of allelic variation governing variables that are relevant for agriculture requires accurate description of alleles in haplotypes (Shi et al. 2019). At present, haplotype-resolved genomes sequencing can fully assemble and annotate the genomes of species with highly heterozygous, and mask allelic variation and functional differentiation of divergent alleles in heterozygous species (Hu et al. 2022; Lin et al. 2022; Han et al. 2023). Finding divergent bialleles on homologous chromosomes is made possible by the high-quality, allele-defined reference genome, which is useful for investigating the differentiation and expression dominance of bialleles as well as the underlying evolutionary processes that underlie these processes (Hu et al. 2021). Our comprehension of the genetic foundation of biallelic variation is enhanced by the haplotype-resolved in oregano, which also makes it possible to strategically take advantage of improvements. Innovative breeding techniques for the Lamiaceae family and other highly heterozygous plants will be made easier by these genomic resources.

Here, two haplotype-resolved *O. vulgare* ’Hot & Spicy’ genomes that were produced by combining chromatin conformation capture (Hi-C) and high-fidelity (HiFi) sequencing technologies (Supplementary Fig. S1). Through haplotype-resolved genomes-scale analyses of the sequencing data, in conjunction with analysis of comparative genomic, phylogenetic, transcriptomic, we desired to illustrate the evolution, mechanism of GSTs formation in oregano. Then, we construced of F_2_ genetic population and high-density genetic linkage maps, identified GST density of leaf adaxial surface (GADS), abaxial surface (GABS), and total surfaces (GTS), and located quantitative trait locus (QTLs) of GADS, GABS, and GTS. Our research will not only provide significant insight into the GSTs formation mechnism, but also facilitate molecular breeding in oregano.

## Results

### Phenotypic evaluation of two oregano

We evaluated two oregano of *O. vulgare* ‘Hot & Spicy’ and *O. vulgare*, one of which is European cultivated variety from the Czech Republic, the other one is representative Chinese wild species. The EO yield (extraction rate), GSTs type and density, EO compositions and their contents are shown in Supplementary Fig. S1 and Fig. 4B. EO yields of *O. vulgare* ‘Hot & Spicy’ and *O. vulgare* are 2.80% and 0.12%, respectively. The type and size of GSTs are shown in Fig. 4B, including two types of GSTs in oregano, namely CGT and peltate glandular trichome (PGT). The density of GSTs on different tissues in *O. vulgare* ‘Hot & Spicy’ is significantly higher than that in *O. vulgare* (Fig. 4A). For example, the densities of PGTs on adaxial and abaxial leaves surfaces (per cm^2^) of *O. vulgare* ‘Hot & Spicy’ and *O. vulgare* are 500/400 and 42/153, respectively. EO compositions and their contents are shown in Supplementary Fig. S1. Carvacrol (89.90%) is the main aromatic composition in *O. vulgare* ‘Hot & Spicy’, whereas caryophyllene oxide (37.96%) and caryophyllen (16.76%) are the main aromatic compositions in *O. vulgare*. Carvacrol is monoterpene, whereas caryophyllene oxide and caryophyllen are sesquiterpenes. One F_2_ hybrid population was designed using *O. vulgare* ‘Hot & Spicy’ as the female parent and *O. vulgare* as the male parent.

### Sequencing and assembly of oregano haplotype-resolved genomes

We sequenced two haplotype-resolved genomes of *O. vulgare* ‘Hot & Spicy’ (2n = 2x = 30). The two haplotype-resolved genomes size of *O. vulgare* ‘Hot & Spicy’ were estimated to be both 502.52 Mb based on k-mer counting, respectively (Supplementary Data Set 1; Supplementary Fig. S2). The k-mer distribution analysis (k = 17) revealed a primary peak at 36.8×, suggesting ∼2.83% high level of heterozygosity and 71.86% highly repetitive sequence content in the genomes, respectively. To obtain two haplotype-resolved genomes, 49.62 gigabases (Gb) of PacBio circular consensus sequencing (CCS) reads were generated using the PacBio Sequel II platform (Supplementary Data Set 1). Contigs were corrected and scaffolded using Hi-C into all 15 pseudochromosomes of two haplotype-resolved genomes (Supplementary Data Set 1; Supplementary Figs. S3–S8). Finally, two haplotype-resolved genomes were obtained, which contained 15 pseudochromosomes with the total length of 606.75 Mb and 612.74 Mb, with the contig N50 of 23.22 Mb and 32.52 Mb, and BUSCO of 94.92% and 94.98%, respectively (Supplementary Data Set 1; Fig. 2; Supplementary Data Sets 2–6; Supplementary Figs. S9 and S10).

**Figure 1.**
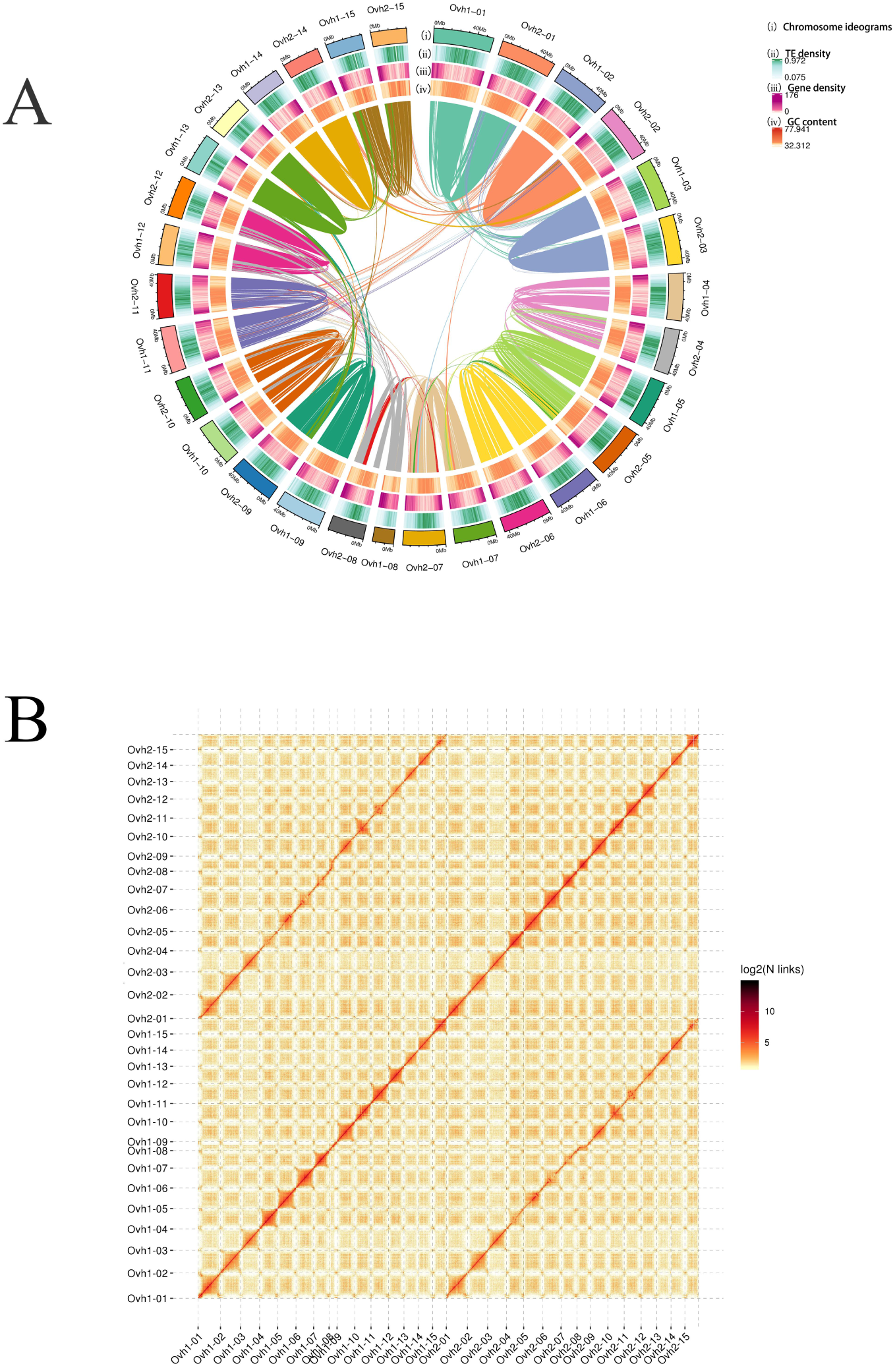
Overview of two haplotype-resolved genomes of *O. vulgare* ‘Hot & Spicy’. **A)** Oregano genomic features. i Chromosome ideograms; ii TE density; iii Gene density; iv GC content. **B)** Heatmap of genomic interactions of two haplotype-resolved genomes among 15 pseudochromosomes. Schematic presentation of major inter-chromosomal relationships in the *O. vulgare* ‘Hot & Spicy’ hap1 (Ovh1-01– Ovh1-15) and *O. vulgare* ‘Hot & Spicy’ hap2 (Ovh2-01–Ovh2-15) genomes. The chromosome size is shown in Mb. Mb, megabases.

**Figure 2.**
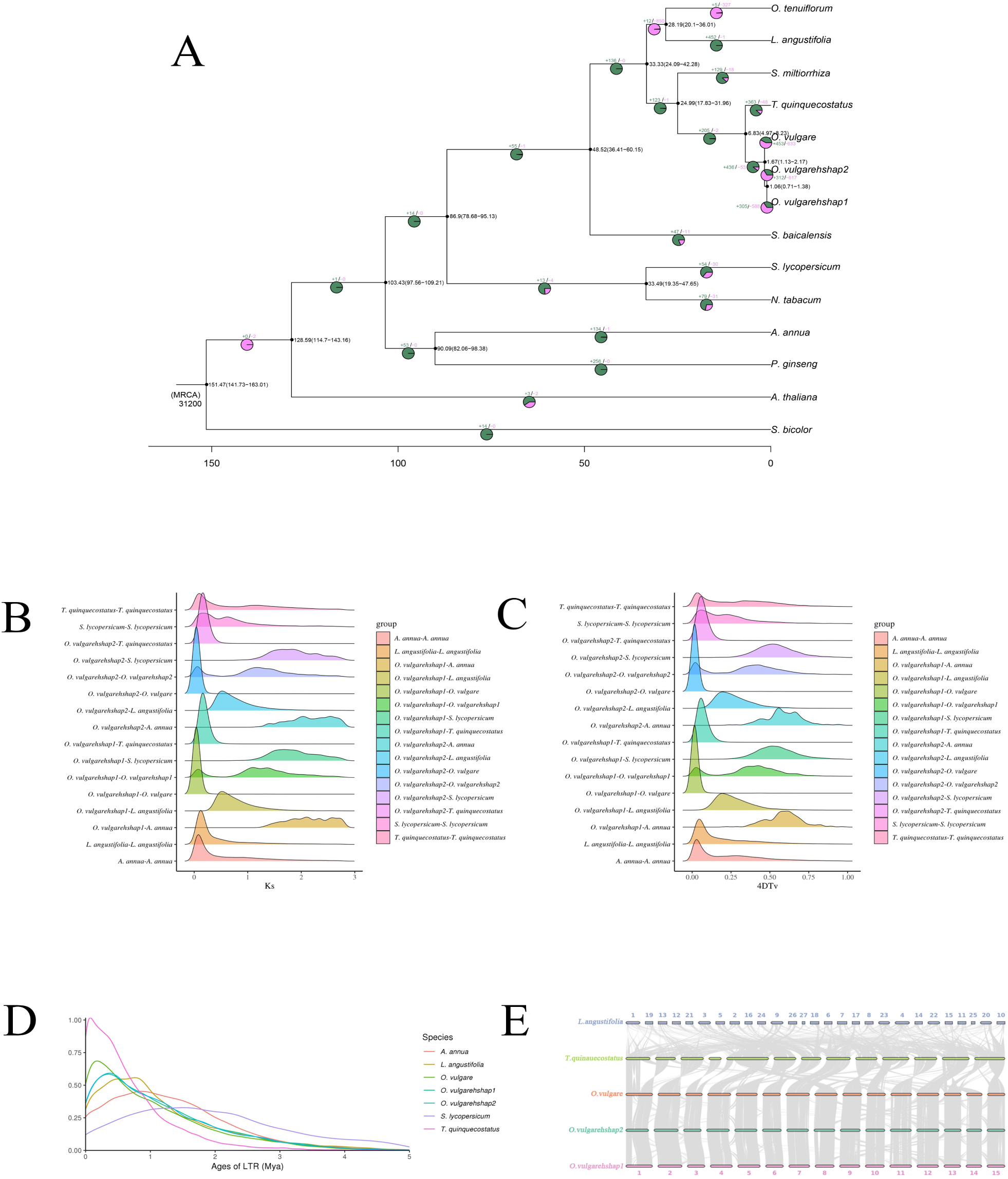
Evolutionary, comparative genomic, and synteny analyses of the two haplotype-resolved genomes. **A)** Phylogenetic analysis, divergence time estimations, and the number of expansion and contraction of gene families and their distribution among 13 plant species. The tree was constructed based on 1,634 single-copy truly orthologous genes. Divergence times (Mya) are indicated by the red numbers beside the branch nodes. The number of gene-family expansion and contraction events is indicated by green and pink numbers (respectively) on each species branch. MRCA, the most recent common ancestor; Mya, million years ago. **B)** Distribution of the synonymous substitution rate (Ks) between *A. annua*, *L. angustifolia*, *O. vulgare*, *O. vulgare hshap1*, *O. vulgare hshap2*, *S. lycopersicum*, and *T. quinquecostatus*, respectively. **C)** Genome duplication in *A. annua*, *L. angustifolia*, *O. vulgare*, *O. vulgare hshap1*, *O. vulgare hshap2*, *S. lycopersicum*, and *T. quinquecostatus* genomes as revealed through 4DTv analyses. **D)** Distribution of insertion ages of LTR-retrotransposons in the genomes of *O. vulgare hshap1*, *O. vulgare hshap2*, and related 12 species genomes. LTR, long terminal repeat. **E)** Visualization of collinear blocks between *O. vulgare hshap1*, *O. vulgare hshap2*, *O. vulgare*, *T. quinquecostatus*, and *L. angustifolia*.

Benchmarking Universal Single-Copy Orthologs (BUSCO), Core Eukaryotic Genes Mapping Approach (CEGMA), and short-read sequence alignment were used to verify the completeness and quality of the produced two haplotype-resolved genomes. 96.08% (245 of 255 core eukaryotic genes) and 96.86% (247 of 255 core eukaryotic genes) of the total genes in two haplotype-resolved genomes, respectively. The conserved genes belonging to complete and single-copy, complete and duplicated, fragmented, and missing categories accounted for 225 (88.24%)/225 (88.24%), 20 (7.84%)/22 (8.63%), 5 (1.96%)/2 (0.78%), and 5 (1.96%)/6 (2.35%) of the total genes.

By searching conserved eukaryotic genes in the assembly using CEGMA, the outputs showed that 97.60%/97.60% of the eukaryotic conserved genes (447/447 of 458 eukaryotic conserved genes) and 96.37%/96.77% of the core eukaryotic genes (239/240 of 248 highly conserved eukaryotic genes) were present in two haplotype-resolved genomes hap1 and hap2, respectively.

The pseudochromosomes assembly were obtained using ALLHiC with 140.17 Gb of Hi-C clean data (Supplementary Data Sets 4 and 5) in *O. vulgare* ‘Hot & Spicy’. A total of 150 and 189 contigs were constructed with the contig N50 of 23.22 and 32.52 Mb, and the longest contig of 45.87 and 46.52 Mb, respectively (Supplementary Data Set 1). A total of 131 and 174 scaffolds were constructed with the scaffold N50 of 42.95 and 41.81 Mb, and the longest scaffold of 54.44 and 51.01 Mb, respectively (Supplementary Data Set 1). A total of 606.75 and 612.74 Mb of sequences were anchored onto 15 pseudochromosomes, accounting for 97.42% and 96.98% of the initial assembly in two haplotype-resolved genomes hap1 and hap2, respectively (Supplementary Data Set 1). In addition, Hi-C data were mapped against the Hi-C scaffold assembly, showing 602.49 and 611.22 Mb of sequences for determining the order and direction, and the 99.30% and 99.75% valid rate of assembled sequences, respectively (Supplementary Data Sets 5 and 6).

### Genome annotation and repeat elements in two oregano genomes

A total of 28,931 and 29,173 protein-coding genes were predicted, and 28,491 (99.05%) and 28,772 (98.57%) were annotated by incorporating transcriptome, homology, and abinitio prediction in two haplotype-resolved genomes hap1 (*O. vulgare hshap1*) and hap2 (*O. vulgare hshap2*), respectively (Supplementary Data Set 7). Statistical gene information of two haplotype-resolved genomes, and related species *Arabidopsis thaliana*, *Lavandula angustifolia*, *Ocimum tenuiflorum*, *O. vulgare* (Sun et al. unpublished data)*, Scutellaria baicalensis*, *Salvia miltiorrhiza*, and *Thymus quinquecostatus* showed the average gene length of 3,487.13 and 3,501.48 bp, the coding sequence length of 1,813.79 and 1,830.11 bp, and the exon number of 5.26 and 5.36, respectively ( Supplementary Data Set 8; Supplementary Figs. S3 and S4).

We functionally annotated these genes against published databases, including Pfam, Swissprot, TrEMBL, eggNOG (Supplementary Figs. S5A and S6A), and nr, resulting in 88.16%/88.53%, 78.62%/78.29%, 98.29%/98.43%, 85.42%/85.95%, and 96.56/96.68% functionally assigned genes in two haplotype-resolved genomes hap1 and hap2, respectively (Supplementary Data Set 9). We further annotated these genes using Eukaryotic Orthologous Groups (KOG), Gene Ontology (GO) (Supplementary Figs. S5B and S6B), and Kyoto Encyclopedia of Genes and Genomes (KEGG) databases. Approximately 56.83% and 57.42% of genes showed orthologous groups in KOG, 82.64% and 82.92% showed GO term classification, and 75.46% and 75.77% could be mapped to known plant biological pathways in two haplotype-resolved genomes hap1 and hap2, respectively (Supplementary Data Set 9).

Two haplotype-resolved genomes were highly repetitive with the total of 433.42 and 443.35 Mb of repetitive sequences annotated, accounting for 69.60% and 70.17% in two haplotype-resolved genomes hap1 and hap2, respectively (Supplementary Data Set 1). Long terminal repeat (LTR) retrotransposons were the dominant repeat type, taking up 411.78 (66.12%) and 421.14 (66.65%) Mb of the two haplotype-resolved genomes, respectively (Supplementary Data Set 10 and 11). The LTRs consist of two major types as Class I (Retroelement) and class II (DNA transposon) representing 375.65 (60.32%)/383.31 (60.67%) Mb and 36.13 (5.80%)/37.83 (5.99%) Mb of the two haplotype-resolved genomes, respectively. In addition, a total of 21,651,483 and 22,211,219 bp tandem repeats were identified, accounting for 3.48% and 3.52% Mb of the two haplotype-resolved genomes, respectively (Supplementary Data Set 12). We further identified 3,336/2,860 rRNAs, 738/763 tRNAs, 33/32 micro RNAs (miRNAs), 120/115 small nuclear RNAs (snRNAs), 169/164 small nucleolar RNAs (snoRNAs), and 373/352 pseudogenes in two haplotype-resolved genomes, respectively (Supplementary Data Set 1).

### Phylogenetic and comparative genomic analyses of two haplotype-resolved genomes

To study the evolutionary history and divergence of oregano, we performed comparative genomic analyses of *O. vulgare* ‘Hot & Spicy’ with the genomes of the 12 selected angiosperm species, including *Artemisia annua*, *A. thaliana*, *L. angustifolia*, *Nicotiana tabacum*, *O. tenuiflorum*, *O. vulgare* (Sun et al. unpublished data), *Panax ginseng*, *S. baicalensis*, *S. miltiorrhiza*, *Solanum lycopersicum*, *Sorghum bicolor*, and *T. quinquecostatus*, respectively (Supplementary Data Set 13). The maximum-likelihood-based phylogenetic analyses were performed with *S. bicolor* as the outgroup. The most recent common ancestor (MRCA) of 13 aforementioned species contained 50,202 gene families and 1,634 high-quality single-copy orthologous genes (Fig. 2). Molecular dating using *S. bicolor* as the fossil calibration indicated that two haplotype-resolved genomes hap1 and hap2 emerged ∼1.06 million years ago (Mya), *O. vulgare* and *O. vulgare* ‘Hot & Spicy’ emerged ∼1.67 Mya, and *Origanum*-*Thymus* emerged ∼6.83 Mya.

A total of 50,202 gene families were identified from 13 selected angiosperm species, and 1,808 common gene families were identified in the genomes (Supplementary Data Set 14; Supplementary Figs. S11 and S12). Compared to the other 12 angiosperm plants, two haplotype-resolved genomes hap1 and hap2 showed 52 and 75 unique gene families (Supplementary Fig. S12A). We compared the gene families among five Lamiaceae species which showed 13,880 gene families were shared by *O. vulgare hshap1*, *O. vulgare hshap2*, *O. vulgare*, *T. quinquecostatus*, *L. angustifolia*, and *S. baicalensis*, and 125 and 151 gene families were specific to two haplotype-resolved genomes hap1 and hap2, respectively (Supplementary Fig. S13). Comparative analyses of contraction, expansion, and positive selection gene families in 13 plants (Fig. 2A) showed that 2,517/2,726 genes had expanded, 626/529 had contracted, and 149/124 had positive selected in two haplotype-resolved genomes hap1 and hap2, respectively (Supplementary Figs. S14–S16). The functional annotation of contraction, expansion, and positive selection gene families accounts for various traits of two haplotype-resolved genomes, including high level of terpenes.

### Whole-genome duplication and synteny analyses of two haplotype-resolved genomes

The analysis of the synonymous substitution rate (Ks) revealed a peak Ks distribution of ∼0.025 for *O. vulgare hshap1*-*O. vulgare,* ∼0.024 for *O. vulgare hshap2*-*O. vulgare*, ∼0.142 for *O. vulgare hshap1*-*T. quinquecostatus,* ∼0.150 for *O. vulgare hshap2*-*T. quinquecostatus*, ∼0.500 for *O. vulgare hshap1*-*L. angustifolia,* and ∼0.504 for *O. vulgare hshap2*-*L. angustifolia*, corresponding to the divergence ∼1.435, ∼1.430, ∼8.294, ∼8.757, ∼29.124, and ∼29.470 Mya, respectively (Fig. 2B). Two whole-genome duplication (WGD) events were investigated in the *O. vulgare hshap1* and *O. vulgare hshap2* genomes based on the distribution of Ks values ∼0.088/0.078 and ∼1.143/1.196 between orthologs, corresponding to the divergence ∼5.120/4.564 and ∼66.857/69.923 Mya, respectively.

Further analysis of the four-fold degenerative synonymous third-codon transversion (4DTv) values for syntenic duplicate genes revealed two significant peaks for two haplotype-resolved genomes *O. vulgare hshap1* and *O. vulgare hshap2* (4DTv ∼0.031/0.029 and ∼0.409/0.411; Fig. 2C), which further confirmed that two oregano genomes had experienced two WGD events. A divergence peak value (4DTv ∼0.0114 and ∼0.0111) was observed for *O. vulgare hshap1*-*O. vulgare* and *O. vulgare hshap2*-*O. vulgare* in the map (Fig. 2C), which suggested that the divergence of *O. vulgare hshap2*-*O. vulgare* occurred later than that of *O. vulgare hshap1*-*O. vulgare*. We found that two wild oregano, like many other flowering plants, had experienced two rounds of WGD with the recent one occurring around ∼5.120 and ∼4.564 Mya followed by extensive genomic rearrangements that resulted in 15 pseudochromosomes in oregano since they diverged from their common paleopolyploid ancestor (Wei et al. 2018; Xia et al. 2020).

Accumulation of LTR-retrotransposons (LTR-RTs) is an important contributor to genome expansion and diversity (Kidwell et al. 1997). The comparison of the insertion ages for LTR-RTs showed similar insertion profiles among the genomes of 13 selected angiosperm species (Fig. 2D). We found that most LTR-RTs insertion events in the two haplotype-resolved genomes *O. vulgare hshap1* and *O. vulgare hshap2* occurred recently or less than 1 Mya. Nevertheless, we also observed that the genomes of two haplotype-resolved genomes *O. vulgare hshap1* and *O. vulgare hshap2* carried younger LTR-RTs, with insertion times 0.35 and 0.37 Mya, respectively (Fig. 2D).

A total of 546, 449, 642, and 2,392 syntenic blocks were identified based on the orthologous gene orders between *O. vulgare hshap1*-*O. vulgare hshap2*, *O. vulgare hshap2-O. vulgare*, *O. vulgare*-*T. quinquecostatus*, and *T. quinquecostatus*-*L. angustifolia* corresponding to 26,084, 24,737, 24,722, and 46,480 gene pairs in *O. vulgare hshap1-O. vulgare hshap2, O. vulgare hshap2-O. vulgare, O. vulgare-T. quinquecostatus, and T. quinquecostatus-L. angustifolia*, respectively (Fig. 2E). A total of 791/1023 syntenic blocks and 7,113/9,050 gene pairs were identified in the inter-genomic comparisons of *O. vulgare hshap1-O. vulgare hshap1* and *O. vulgare hshap2-O. vulgare hshap2*, respectively (Fig. 2E). *Origanum*-*Thymus* had a higher collinearity than *Origanum*-*Lavandula*, as seen by the dot and bar graphs. This finding was in keeping with their close phylogenetic relationship as Lamiaceae clade members.

### Structural variation among two haplotype-resolved genomes

Comparison of two haplotype-resolved genomes hap1 and hap2, SNPs, insertions and deletions (InDels) were detected by SyRI and annotated using the ANNOVAR software toolkit (Fig. 3A). The total number of SNPs was 2,669,410, with in gene region was 960,035 and intergenic region was 1,709,375. we found there were 24,400, 34,905, 140,269, 184,223, 44,664, 319,491, 211,737, 1,709,375, and 346 SNPs in UTR5, UTR3, downstream and upstream of gene region with 1 kb, both upstream and downstream of gene region, intronic, exonic, intergenic, and splicing region with 2 bp. The total number of InDels was 569,093, with 355,543 insertions and 213,550 deletions. we found there were 12,613, 12,375, 49,052, 67,603, 19,290, 91,520, 9,486, 306,843, and 311 SNPs in UTR5, UTR3, downstream and upstream of gene region with 1 kb, both upstream and downstream of gene region, intronic, exonic, intergenic, and splicing region with 2 bp. The intragenic SNPs and InDels were clustered by KEGG analysis in purine metabolism, ubiquitin mediated proteolysis, terpenoid backbone biosynthesis, isoflavonoid biosynthesis, galactose metabolism, ubiquinone and other terpenoid-quinone biosynthesis, etc (Supplementary Fig. S17).

**Figure 3.**
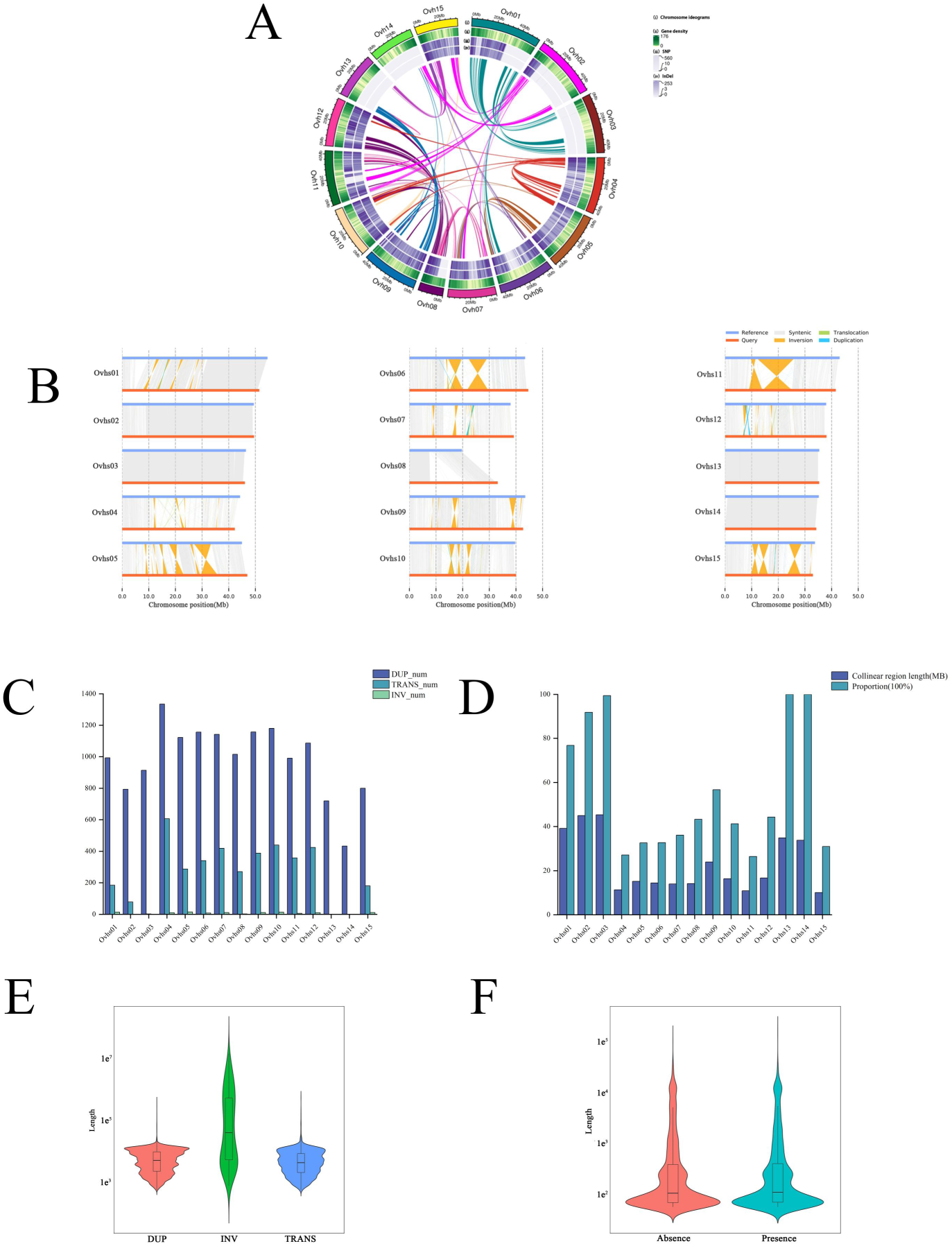
Large number of variations among oregano two haplotype-resolved genomes. **A)** Variation detection of single nucleotide polymorphism (SNP), and insertion and deletion (InDel) among oregano two haplotype-resolved genomes. i Chromosome ideograms; ii Gene density; iii SNP; iv InDel. **B)** Syntenic analyses among oregano two haplotype-resolved genomes. Syntenic regions and structure variantions (SVs) (Inversion, Translocation, and Duplication) are highlighted by different colors. **C)** The total number of duplications (DUPs), inversions (INVs), and translocations (TRANSs) in the 15 pseudochromosomes. **D)** The length and coverage proportion of syntenic regions on the 15 pseudochromosomes. **E)** The length distribution of DUPs, INVs, and TRANSs among oregano two haplotype-resolved genomes. **F)** The length distribution of absences and presences among oregano two haplotype-resolved genomes.

The total number of structure variantions (SVs) was 18,925, with 14,839 duplications (DUPs), 110 inversions (INVs), and 3,976 translocations (TRANSs) in the 15 pseudochromosomes (Fig. 3B). In 15 pseudochromosomes, the difference of each SVs was significant that the numbers of SVs included 433–1,335 DUPs, 0–15 INVs, and 0–606 TRANSs (Fig. 3C). The largest number of SVs were 1335 DUPs in Ovhs04, 15 INVs in Ovhs05, and 606 TRANSs in Ovhs04. The coverage of syntenic regions on the 15 pseudochromosomes ranges from 27.10% (Ovhs04) to 100% (Ovhs13 and Ovhs14)(Fig. 3D). The length distribution of DUPs, INVs, and TRANSs were 501–385,357 bp, 684–14,457,258 bp, and 507–538141 bp, respectively (Fig. 3E). The SVs were clustered by KEGG analysis in purine metabolism, ubiquinone and other terpenoid-quinone biosynthesis, arginine and proline metabolism, linoleic acid metabolism, biosynthesis of unsaturated fatty acids, etc (Supplementary Fig. S18). The total number of presence-absence variations (PAV) was 5,425 with 2,819 absences and 2,606 presences. The length distribution of absences and presences were 50–99,485 bp and 50–149,642 bp, respectively (Fig. 3F). The total number of copy number variation (CNV) was 1,426, with 746 copy gains (CPGs) and copy losses (CPLs) in hap2 comparing hap1. These large structural variations contributed to significant differences in oregano phenotypes.

### RNA-seq analysis reveals the genetic mechanism of GSTs formation

To investigate the genes that play important roles in GSTs formation, we performed transcriptomic analyses of *O. vulgare* ‘Hot & Spicy’ and *O. vulgare* using four different tissues (root, stem, leaf, and flower). A total of 24 cDNA libraries were processed for transcriptome sequencing, generating 147.25 Gb of clean data (Supplementary Fig. S19). At least 5.72 Gb of clean data were generated for each sample with a minimum of 92.99% of clean data having a quality score of Q30 (Supplementary Data Set 15). Clean reads of each sample were mapped to the two haplotype-resolved genomes hap1 and hap2. The mapping ratio reads ranged 86.09– 91.03% and 86.09–91.03%, respectively (Supplementary Data Set 16).

GSTs can synthesize, store, or secrete terpenoids. The type, size, and density of GSTs are shown in Fig. 4. The density of GSTs on flowers and leaves in *O. vulgare* ‘Hot & Spicy’ were significantly higher than those in *O. vulgare* (Fig. 4A). There are two types of GSTs in oregano, which include CGT and PGT (Fig. 4B). By searching and collecting regulatory genes related to GSTs formation in *A. annua*, tobacco, and tomato, we summed up the genetic network of GSTs formation in oregano (Fig. 4C). There are four steps in the creation of GSTs: morphogenesis, maturation, initiation, and determination. It is hypothesized that some GSTs have comparable developmental events due to their shared organizational structure. The transcription factors R2R3-MYB and HD-ZIP IV control the start of most GSTs (TF).

**Figure 4.**
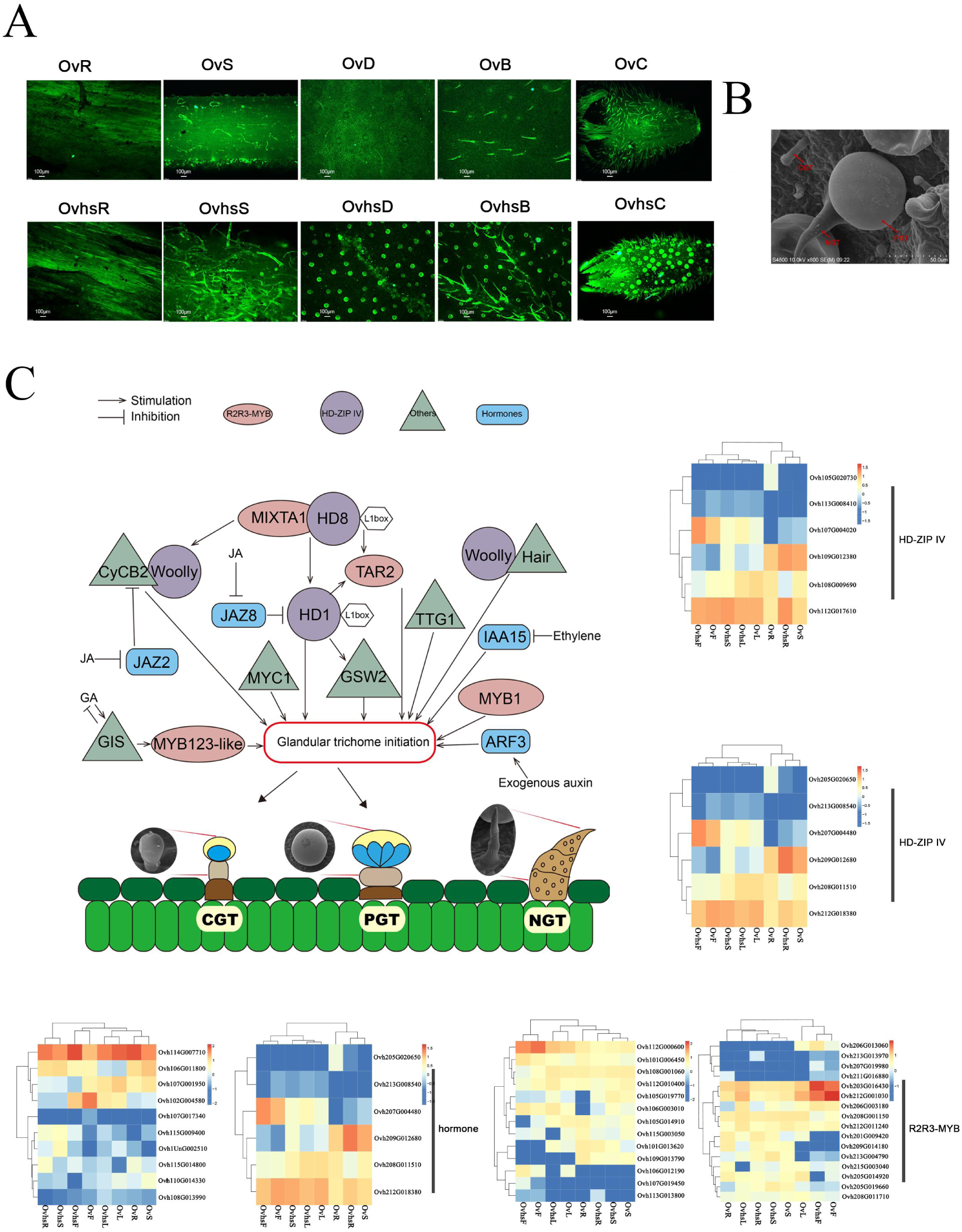
Schematic overview of GST formation mechanism in oregano. **A)** Density of GSTs in different tissues are shown in *O. vulgare* and *O. vulgare* ‘Hot & Spicy’ using fluorescence microscope. OvR, *O. vulgare* root; OvS, *O. vulgare* stem; OvD, *O. vulgare* adaxial leaf; OvB, *O. vulgare* abaxial leaf; OvC, *O. vulgare* calyx; OvhsR, *O. vulgare* ‘Hot & Spicy’ root; OvhsS, *O. vulgare* ‘Hot & Spicy’ stem; OvhsD, *O. vulgare* ‘Hot & Spicy’ adaxial leaf; OvhsB, *O. vulgare* ‘Hot & Spicy’ abaxial leaf; OvhsC, *O. vulgare* ‘Hot & Spicy’ calyx. Bars = 100 μm. **B)** The type of GSTs. CGT, capitate glandular trichome; PGT, peltate glandular trichome; NGT, non-glandular trichomes; GST, glandular secretory trichome. **C)** Expression pathway and heat maps of GST formation mechanism in oregano. Red denotes high expression and blue indicates low expression, expression data are Z-score standardized to -3 to 3 per gene. All of expression genes showing minimum two-fold changes at p < 0.05 (Student’s two tailed t-test).

Based on reported TFs, we summed up the genetic network of GST formation in oregano (Fig. 4C). The process is mainly involved in two important transcription factors including R2R3-MYB (encoded by *MYB1*, *MIXTA1*, *TAR2,* and *MYB123*-like) and HD-ZIP IV (encoded by *HD8*, *HD1,* and *Woolly*). Besides, hormone-related genes (*IAA15*, *JAZ2*, *JAZ8* and *ARF3*) and other genes, such as *GSW2*, *MYC1*, *CyCB2*, *GIS*, *TTG1* and *Hair,* are also involved in this process. The heatmap showed that 13/16 differentially expressed R2R3-MYB-coding genes and 6/6 differentially expressed HD-ZIP IV-encoding genes were involved in GSTs formation two haplotype-resolved genomes hap1 and hap2, respectively (Fig. 4C). We also identified 10/11 hormone-related genes (4/3 *IAA15*, 3/3 *JAZ2*, 1/3 *JAZ8*, and 2/2 *ARF3*) and the other genes, such as 5/4 *GSW2s*, 5/5 *MYC1s*, 3/3 *CyCB2*, 4/4 *GIS*, 3/3 *TTG1,* and 2/2 *Hair* differentially expressed in two haplotype-resolved genomes hap1 and hap2, respectively (Fig. 4C). The interpretation here is that the regulation of these trichome-related genes may underlie the regulation of GSTs density to increase the monoterpenoid and sesquiterpenes content.

### F_2_ genetic population re-sequencing and construction of high density genetic linkage maps

Two parents *O. vulgare* ‘Hot & Spicy’ and *O. vulgare* were sequenced at a depth of 46.50 and 37.18 Gb, respectively (Supplementary Data Set 17). A total of 778.81 Gb pairs of sequencing data was obtained for 232 F_2_ progeny individuals from the cross, with an average of 3.36 Gb pairs per sample (Supplementary Data Set 18). This corresponds to ∼4.42-fold and ∼4.38-fold coverage of two haplotype-resolved genomes hap1 and hap2 for each of the progeny, respectively. Alignment of cleaned data to two haplotype-resolved genomes hap1 and hap2 produced an average mapping rate of 93.82% and 93.72%, respectively (Supplementary Data Sets 19 and 20; Supplementary Fig. S20). Variant calling identified all SNPs and InDels between two parents comparing two haplotype-resolved genomes hap1 or hap2, respectively (Supplementary Data Sets 21–S24; Supplementary Fig. S21). A total of 3,083,292 or 3,296,483 SNPs were detected between parents, and Ti(Transition)/Tv(Transversion) was 1.81 or 1.83, comparing two haplotype-resolved genomes hap1 or hap2, respectively. In order to ensure the quality of the markers in the genetic linkage maps of F_2_ genetic population, the SNP markers were classified and screened by Binmarker v2.3. Finally, 7,325 (61,799 SNPs) and 7,574 (64,571 SNPs) filtered bin markers were obtained which could be used for genetic map analysis, respectively (Supplementary Data Sets 25 and 26). We obtained 5,643 and 6,235 bin markers by illustrated genotype analysis using HighMap software, respectively (Supplementary Figs. S22 and S23).

The resulting hap1 and hap2 genetic linkage maps consisted of 15 linkage groups (LGs) and spanned 2,279.28 and 2,322.83 centimorgan (cM), with an average interval of 0.48 and 0.42 cM (Fig. 5). The max gap distance ranged from 1.77 (LGh1-14) to 8.22 cM (LGh1-10) of hap1 linkage map, meanwhile the max gap distance ranged from 2.20 (LGh2-14) to 8.22 cM (LGh1-10) of hap2 linkage map (Supplementary Data Sets 27 and 28). The genetic length of the LGs ranged from 113.86 (LGh1-09) to 179.63 cM (LGh1-04) with an average length of 151.95 cM of hap1 linkage map (Fig. 5A; Supplementary Data Set 27). There were on average 376.20 bin markers per hap1 linkage group. LGh1-02 contained the fewest bin markers (n = 186) and had the widest mean interval (0.87 cM) between adjacent bin markers in hap1 linkage map. LGh1-02 contained highest number of bin markers (n = 681) and the narrowest mean interval in hap1 linkage map (0.21 cM). The genetic length of the LGs ranged from 120.28 (LGh2-10) to 199.31 cM (LGh2-07) with an average length of 154.86 cM of hap2 linkage map (Fig. 5B; Supplementary Data Set 28). There were on average 415.67 bin markers per hap2 linkage group. LGh2-10 contained the fewest bin markers (n = 226) and had the widest mean interval (0.77 cM) between adjacent bin markers in hap1 linkage map. LGh2-15 contained highest number of bin markers (n = 753) and the narrowest mean interval in hap1 linkage map (0.23 cM).

**Figure 5.**
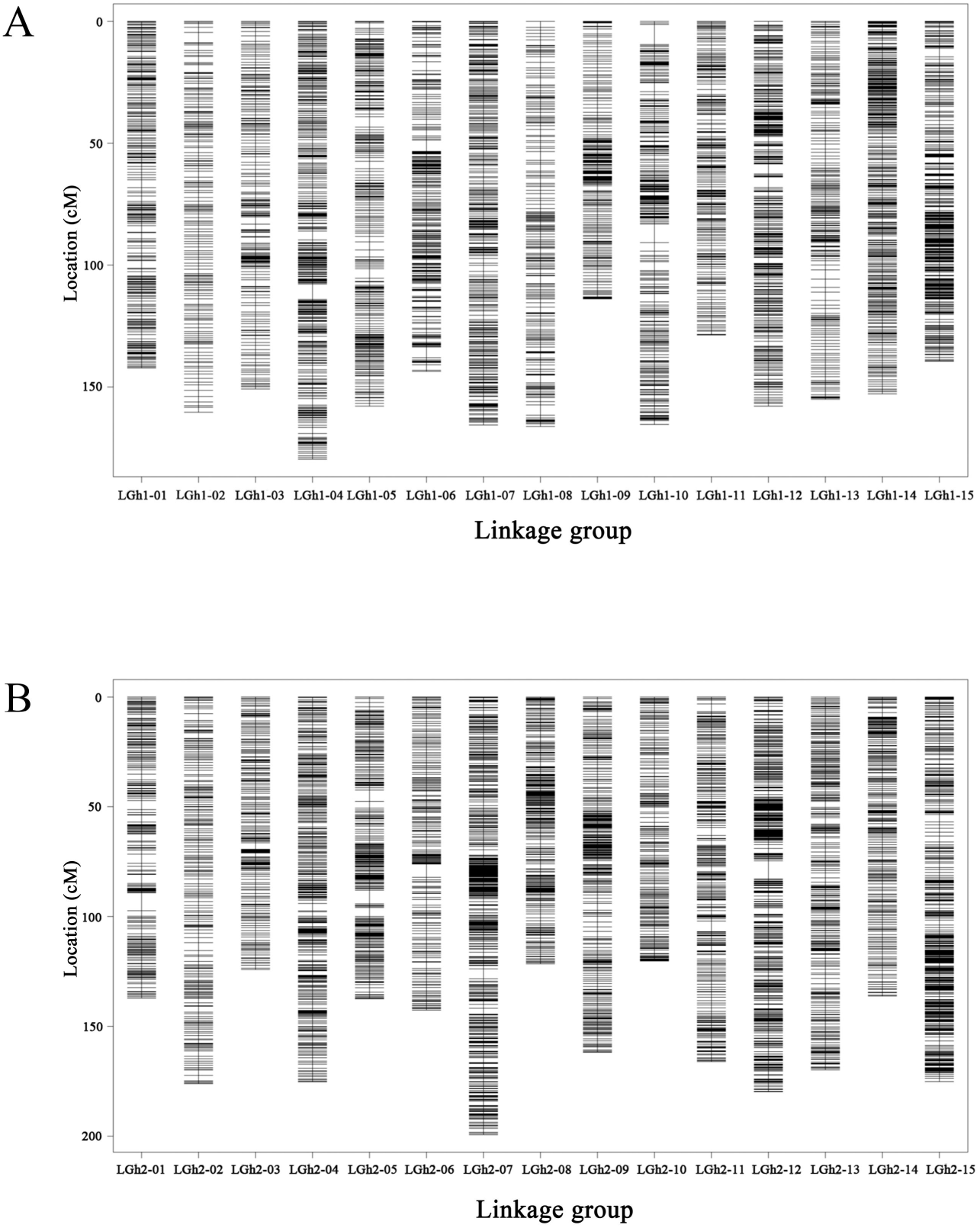
High-density genetic map hap1 and hap2 of F_2_ oregano genetic population based on SNPs. **A)** High-density genetic map hap1 of oregano based on 61,799 SNPs (7,325 bin markers). **B)** High-density genetic map hap1 of oregano based on 64,571 SNPs (7,574 bin markers). The x-axis of each subplot indicates the linkage group (LGh1-01–LGh1-15 and LGh2-01–LGh2-15) and the y-axis indicates the location (cM).

We performed collinearity analysis between physical maps with SNP-based high-density genetic maps to identify SNP-bin pairs located on two haplotype-resolved genomes. The location of the SNP-bin markers on the hap1 and hap2 genomes and the genetic maps were analyzed colinearally which showed in Supplementary Figs. 24–26. Spearman correlation coefficients of physical maps and genetic maps are close to 1 which showed the genetic linkage maps are very well collinearity with the physical maps (Supplementary Data Sets 29 and 30).

### Phenotyping of density of GSTs-related traits and quantitative trait locus analysis

The continuous variation of GST density of GADS, GABS, and GTS traits from the *O. vulgare* ‘Hot & Spicy’ and *O. vulgare* segregating population demonstrated the quantitative nature of these traits (Fig. 6A, B). GADS, GABS, and GTS of female parent *O. vulgare* ‘Hot & Spicy’ were 501.54, 402.36, and 903.90 /cm^2^, meanwhile GADS, GABS, and GTS of male parent *O. vulgare* were 42.55, 153.34, and 195.89/cm^2^. GADS, GABS, and GTS of F_2_ segregating populations were greatly different which showed obvious superparental dominance (Fig. 6A). The GST density of leaf abaxial surfaces were much higher than those on leaf adaxial surfaces. The analysis results showed that three traits GADS, GABS, and GTS had a positive-skewed distributions (Fig. 6B). GADS ranged from 3.54 to 654.34 /cm^2^, GABS ranged from 75.46 to 744.58 /cm^2^, and GTS ranged from 80.93 to 1,251.77 /cm^2^, respectively.

**Figure 6.**
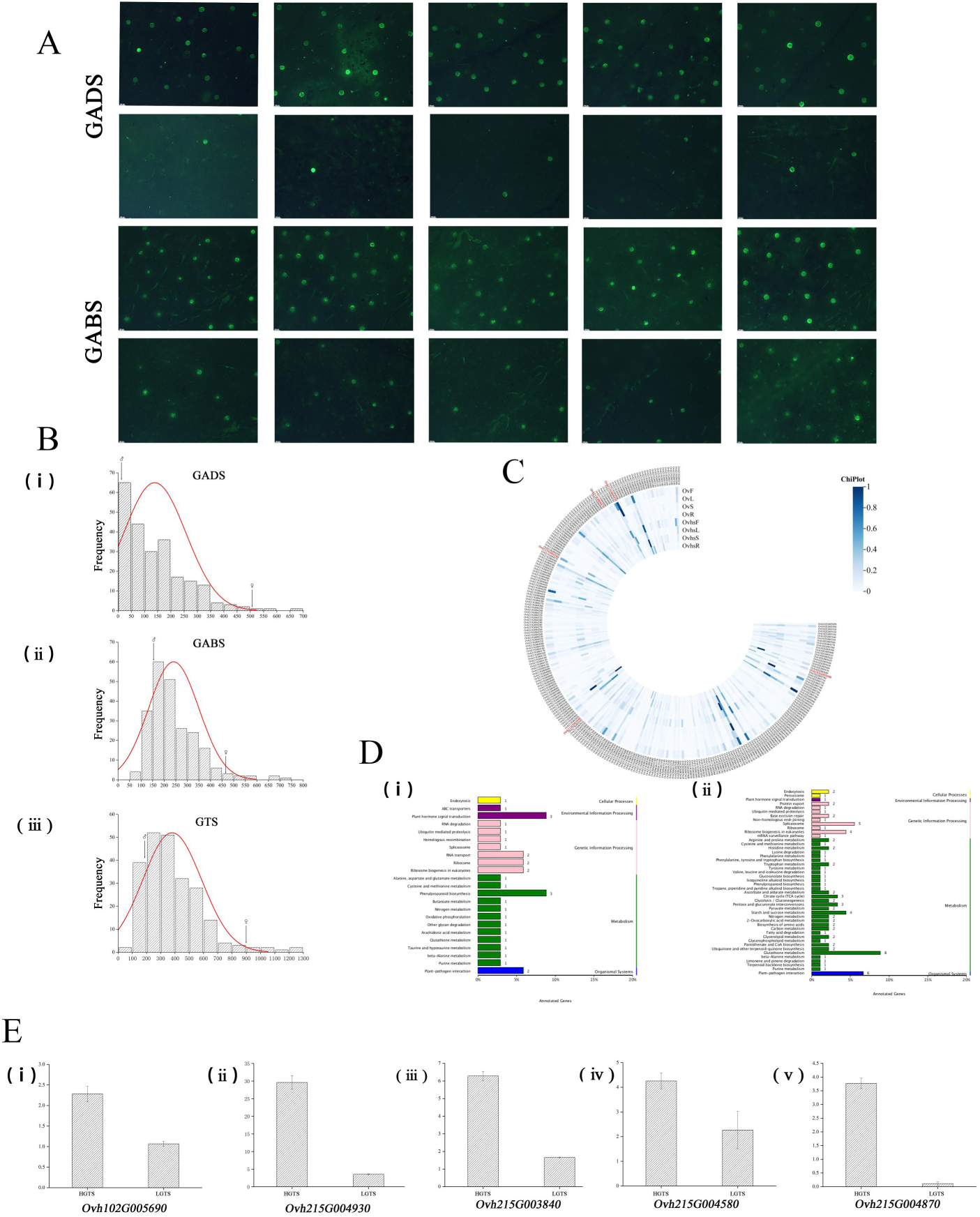
Density of GSTs-related traits phenotyping and candidate gene screening. **A)** Fluorescence microscopy images of the adaxial and abaxial leaf surfaces’ GSTs. GADS, GSTs of leaf adaxial surface; GABS, GSTs of leaf abaxial surface. **B)** Phenotypic distribution of GSTs-related traits in 232 F_2_ progenies from hybridization of *O. vulgare* ‘Hot & Spicy’ and *O. vulgare.* i, GADS, GSTs of leaf adaxial surface; ii, GABS, GSTs of leaf abaxial surface; iii: GTS, GSTs of leaf total surfaces. **C)** Heat map of 367 DEGs in the QTL regions of hap1 and hap2. **D)** KEGG functional annotations for candidate genes in QTL regions of hap1 and hap2. **E)** Candidate genes expression levels in two mixed pools HGTS and HGTS by qRT-PCR. i Expression trends of *Ovh102G005690*; ii Expression trends of *Ovh215G004930*; iii Expression trends of *Ovh215G003840*; iv Expression trends of *Ovh215G004580*; v. Expression trends of *Ovh215G004870*.

In the hap1 linkage map, QTLs for GADS, GABS, and GTS were independently mapped on different linkage groups (Fig. 7A). These QTLs corresponded to GADS on LGh1-15, GABS on LGh1-02, and GTS on LGh1-02 and LGh1-15. The QTL physical regions of GADS on LGh1-15, GABS on LGh1-02, GTS on LGh1-02 and LGh1-15 were 1.26 Mb, 551.56 Kb, 551.56 Kb, and 65.77 Kb. The top region of LGh1-15 (10.17–11.04 cM) showed QTL for GADS; in the middle region of LGh1-02 (20.97–21.40 cM) there were QTL for GABS; and the bottom regions of LGh1-02 (20.97–21.40 cM) and LGh1-15 (7.98–8.41 cM) contained QTLs for GTS. The highest LOD score for GADS was on LGh1-15 (10.21) explaining 18.15% of the phenotypic variation which the additive effect was 81.49 and dominance effect was 8.85. The highest LOD score for GABS was on LGh1-02 (7.30) explaining 9.19% of the phenotypic variation which the additive effect was 57.86 and dominance effect was -17.00. The highest LOD scores for GTS were on LGh1-02 (4.53) and LGh1-15 (4.37) explaining 10.56% and 9.83% of the phenotypic variation, which the additive effect were 116.98 and 103.40, and dominance effect were -24.81 and 16.56, respectively.

**Figure 7.**
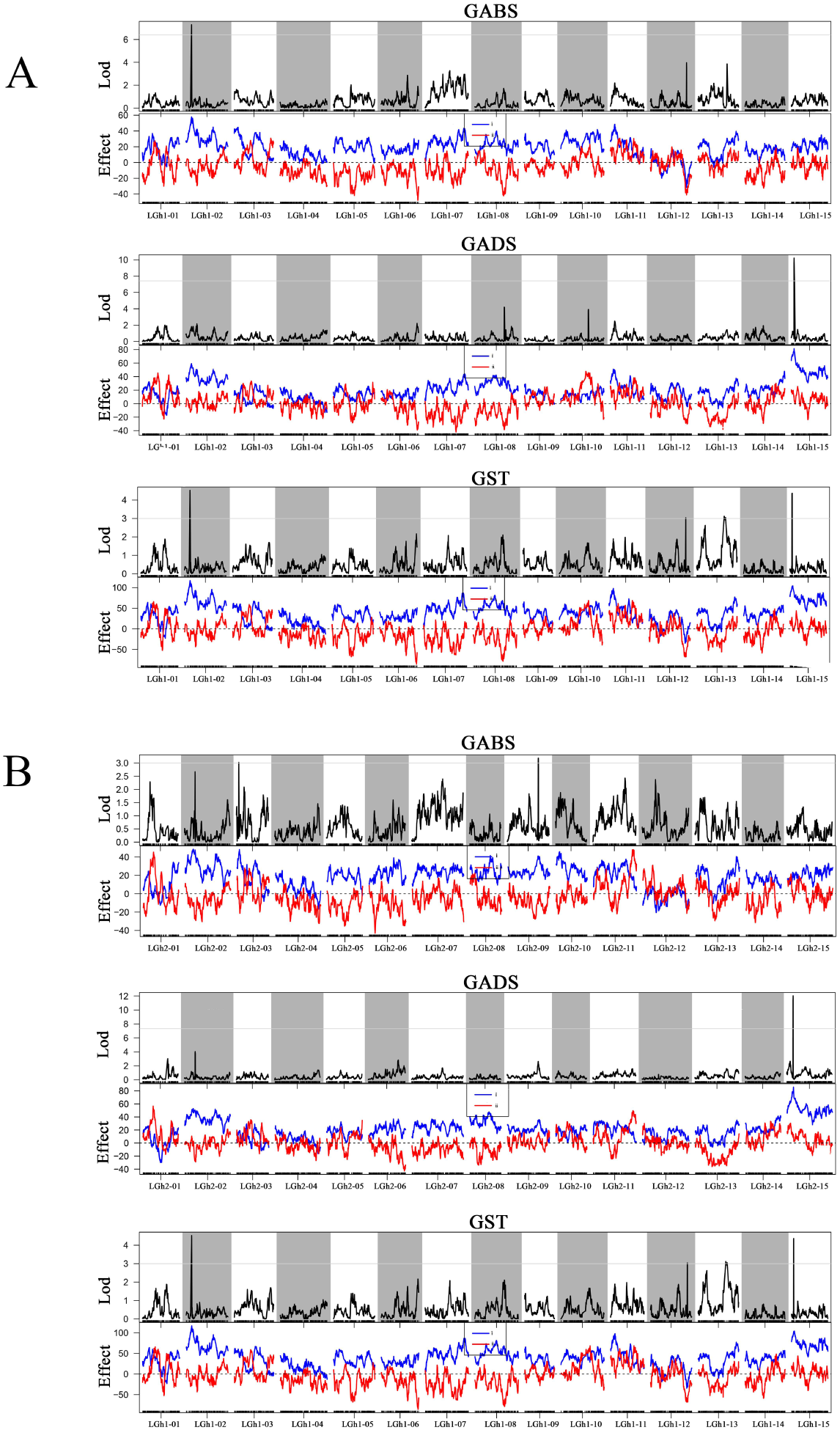
Quantitative trait locus (QTL) analysis of F_2_ oregano genetic population density of GSTs-related traits in genetic map hap1 and hap2. **A)** QTL analysis of F_2_ oregano genetic population density of GSTs-related traits in genetic map hap1. **B)** QTL analysis of F_2_ oregano genetic population density of GSTs-related traits in genetic map hap2. GADS, GSTs of leaf adaxial surface; GABS, GSTs of leaf abaxial surface; GTS, GSTs of leaf total surfaces. The gray lines represent LOD value distribution graphs in the 15 linkage groups. i Blue lines represent additive effects; ii Red lines represent dominance effects.

In the hap2 linkage map, QTLs for GADS, GABS, and GTS were independently mapped on different linkage groups (Fig. 7B). These QTLs corresponded to GADS on LGh2-15, GABS on LGh2-09, and GTS on LGh2-02, LGh2-09, and LGh2-15. The QTL physical regions of GADS on LGh2-15, GABS on LGh2-09, GTS on LGh2-02, LGh2-09, and LGh2-15 were 928.80 Kb, 66.77 Kb, 57.41 Kb, 66.77 Kb, 103.78 Kb and 928.80 Kb. The top region of LGh2-15 (23.88–24.53 cM) showed QTL for GADS; in the middle regions of LGh2-02 (38.12–38.55 cM), LGh2-09 (119.99– 120.21 cM; 143.88–144.52 cM) and LGh2-15 (23.88–24.53 cM) there were QTLs for GTS; and the bottom region of LGh2-09 (119.99–120.21 cM) contained QTL for GABS. The highest LOD score for GADS was on LGh2-15 (12.03) explaining 22.97% of the phenotypic variation which the additive effect was 85.67 and dominance effect was 7.84. The highest LOD score for GABS was on LGh2-09 (3.19) explaining 5.52% of the phenotypic variation which the additive effect was 539.22 and dominance effect was 0.31. The highest LOD scores for GTS were on LGh2-02 (4.10), LGh2-09 (3.22; 3.28) and LGh2-15 (4.07) explaining 10.03%, 5.66%, 1.15%,and 11.57% of the phenotypic variation, which the additive effect were 98.14, 72.62, 26.58, and 107.85, and dominance effect were -54.89, 16.62, -33.77, and 1.72, respectively. In the hap1 and hap2 linkage maps, QTLs for GADS, GABS, and GTS existed differences of the physical and genetic location of pseudochromosomes, phenotypic variation, additive effect, and dominance effect.

The candidate genes associated with the traits were found by using the physical location on two haplotype-resolved genomes hap1 and hap2 corresponding to the bin marker. As a result, a total of 225 (LGh1-15) and 174 (LGh2-15) candidate genes associated screen the candidate genes related to the QTLs of GADS were identified. A total of 79 (LGh1-02) and 2 (LGh2-09) candidate genes associated screen the candidate genes related to the QTLs of GABS were identified. A total of 79 (LGh1-02), 10 (LGh1-15), 6 (LGh2-02), 2 (LGh2-09), 7 (LGh2-09), and 174 (LGh2-15) candidate genes associated screen the candidate genes related to the QTLs of GTS were identified. We performed KEGG gene functional annotations for candidate genes in QTL regions (Fig. 6D). To further to screen candidate genes, we analyzed transcriptome data of two parents at different tissues. Combining transcriptome profiles with the functional annotations, 367 DEGs in the QTL regions of hap1 and hap2 were selected to analyze heat map (Fig. 6C). Subsequently, we combined the DEGs in the QTL regions to detect their expression levels in two mixed pools HGTS and HGTS by qRT-PCR. As shown in Fig. 6E, the expression level of *Ovh102G005690*, *Ovh215G004930*, *Ovh215G003840*, *Ovh215G004580*, and *Ovh215G004870* were relatively high in HGTS, and showed low expression trends in HGTS. We will then verify their genes function by transforming in tomato or tobacco.

## Discussion

The genus *Origanum* is an important aromatic and medicinal plant which has valuable ornamental, flavoring, medicinal, and aromatic properties (Marinelli et al. 2018; Kang et al. 2019; Falleh et al. 2020; Lu et al. 2021; Zhong et al. 2021; Hao et al. 2022; Wang et al. 2023). Oregano EOs contain the important medicinal functional ingredient terpenes such as carvacrol and thymol which have been widely used in folk medicine since ancient times (Marinelli et al. 2018; Hao et al. 2022). The plant EOs compositions have been widely applied as pharmaceuticals, agricultural products, food preservation agent, food additive, feed additive, etc (Kang et al. 2019; Falleh et al. 2020; Lu et al. 2021; Zhong et al. 2021; Wang et al. 2023). In this study, we report two high-quality haplotype-resolved reference genomes for *Origanum* which provide insight into the GSTs formation molecular mechanism. To date, a few of genomes of Lamiaceae have been reported, e.g., *S. miltiorrhiza* (Xu et al. 2016), *S. splendens* (Dong et al. 2018), *S. baicalensis* (Zhao et al. 2019), *Salvia rosmarinus* (Han et al. 2023), *T. quinquecostatus* (Sun et al. 2022), *L. angustifolia* (Li et al. 2021)*, L.* × *intermedia, and L. latifolia* (Li et al. 2023). We now provide two oregano haplotype-resolved genomes sequence, those genomic data will provide a starting point for comparing the genomes of different Lamiaceae members.

At present, haplotype-resolved genomes sequencing can fully assemble and annotate the genomes of species with highly heterozygous, and mask allelic variation and functional differentiation of divergent alleles in heterozygous species (Hu et al. 2022; Lin et al. 2022; Han et al. 2023). Finding divergent bialleles on homologous chromosomes is made possible by the high-quality, allele-defined reference genome, which is useful for investigating the differentiation and expression dominance of bialleles (Hu et al. 2021). Excellent sequences and annotations of haplotype-resolved genomes provide a strong foundation for a variety of genetic investigations, including comparative genomics, evolutionary studies, and deciphering the genomic architecture of desired traits (Varshney et al. 2014). WGD and polyploidization occur more frequently in plant genomes as they evolve (Baek et al. 2018). WGD can cause the retention of duplicate genes, the growth of introns, and the multiplication of repetitive sequences, all of which contribute to the extension of the genome (Schnable et al. 2009). The high heterozygosity of plants may be caused by a higher pace of evolution in highly heterozygous locations. Our comprehension of the genetic foundation of biallelic variation is enhanced by the haplotype-resolved in oregano, which also makes it possible to strategically take advantage of improvements.

Specific chemicals, such as terpenes, alkaloids, and other organic molecules, are produced by specific structures in plants called glandular trichomes. What we know about molecular data pointing to genes about GSTs formation is mainly come from the work on *Artemisia*, tobacco, and tomato (Robert and Tissier 2020) including R2R3-MYB (Lv et al. 2022) and HD-ZIP Ⅳ (Xie et al. 2021) transcription factors, *HOMEODOMAIN PROTEIN 1* (*AaHD1*), *GLANDULAR TRICHOMESPECIFIC WRKY 2* (*AaGSW2*), *JA ZIM-domain 8* (*AaJAZ8*), *WRKY* transcription factor (*AaGSW2*) (Xie et al. 2021), as well as receptors involved in phytohormone-induced signaling cascades (Chen et al. 2023), etc. However, there is less understanding about the GSTs formation mechanism in Lamiaceae species. *A. annua* glandular trichomes are where the vital antimalarial medication artemisinin is biosynthesised and stored, and the density of trichomes is correlated with the amount of artemisinin present (Lv et al. 2022). Increasing the density of GST is one way to improve these compounds’ production. *AaMYB16* was discovered to be a positive regulator of GST initiation by the creation and characterisation of transgenic plants, while *AaMYB5* had the opposite effect (Xie et al. 2021). In this study, we found many transcription factors and genes related to GSTs, such as R2R3-MYB-and HD-ZIP IV-encoding genes in oregano. GST density has been proven to be positively correlated with the production of natural metabolites. Consequently, increasing the density of GSTs would be an effective approach to improve productivity of these natural metabolites (Xie et al. 2021). The implication here is that the modulation of GSTs density to raise the terpene content of oregano may be dependent on the regulation of these trichome-related genes.

Hybridization between or within species can produce heterozygous genomes, which can contribute to the preservation of plant variety and perhaps lead to the emergence of new species (Baek et al. 2018). In agriculture, horticulture, fruit, or tree breeding, it is often possible to construct different populations by hybridization for the generation of new species. Genetic linkage maps have been employed in QTL research to analyze the genetic basis of complex traits in a variety of horticultural and fruit plants, including crape myrtle (Zhou et al. 2023), grape (Zhang et al. 2023), and kiwifruit (Wang et al. 2023), etc. A few studies have identified QTL linked with GST density in plants having GST (Palomares-Rius et al. 2016; Gao et al. 2018; Bennewitz et al. 2018). Using an F4 mapping population and a genetic linkage map created from genotyping-by-sequencing data consisted of 2121 SNP markers, two significant QTL influencing CGT density in sunflower florets were discovered (Gao et al. 2018). A six-generation family and a recombinant inbred lines (F_7_) population were tested for the character in order to ascertain the genetic basis of the density of glandular trichomes type I in melon (Palomares-Rius et al. 2016). The enhanced glandular trichome density resulting from the TGR-1551 alleles for this QTL was validated by confirmation of this marker in advanced backcrosses. *S. habrochaites* LA1777 and *S. lycopersicum* var. *cerasiforme* WVa106 were backcrossed to produce a population of 116 individuals in order to better understand the genetic reasons causing these trichome morphological changes (Bennewitz et al. 2018). It was possible to quantify this attribute by creating a trichome score that takes into account the form of type VI trichomes. In an advanced population produced from one of the back-cross lines, the QTL on chromosome I with the highest LOD score was verified and refined to a 500 gene interval.

Together, we construced of one F_2_ genetic population, two high-density genetic linkage maps, identified QTLs of GADS, GABS, and GTS density. A F_2_ population which included 232 progeny individuals from the cross between *O. vulgare* ‘Hot & Spicy’ and *O. vulgare*. Two genetic linkage maps hap1 and hap2 consisted of 15 LGs (Fig. 5). Through phenotypic identification, we found that there was a huge difference in the density of GADS, GABS, and GTS density. One/one, one/one, and two/four QTLs for GADS, GABS, and GTS were independently mapped on two genetic maps, respectively. Five candidate genes *Ovh102G005690*, *Ovh215G004930*, *Ovh215G003840*, *Ovh215G004580*, and *Ovh215G004870* showed extreme difference in two bulked segregant pools from F_2_ population with highest or lowest GTSs density. In conclusion, the generation of two oregano high-quality haplotype-resolved reference genomes presented here, provides a clear picture of the primary sequence architecture of one European cultivated variety *O. vulgare* ‘Hot & Spicy’.

## Methods

### Plant materials

European cultivated variety *O. vulgare* ‘Hot & Spicy’ was introduced from the Czech Republic (TaxID: 39174). The China wild oregano *O. vulgare* (NCBI: TaxID: 39352) was collected directly from its natural habitat Xinjiang Uygur Autonomous Region Government of China during 2019. To identify and categorize, herbarium specimen was examined by the Institute of Botany, Chinese Academy of Sciences (IB-CAS). It was were grown in an experimental farm by cutting propagation in IB-CAS, Beijing, China. After fresh young leaves were collected from two wild oregano plants, all samples were immediately frozen in liquid nitrogen and stored at -80 °C before DNA extraction.

### Haplotype-resolved genomes of *O. vulgare* ‘Hot & Spicy’ sequencing, assembly and annotation

Haplotype-resolved genomes sequencing, assembly and annotation related methods in Supplementary Method S1.

### Variation detection

Genome assemblies were aligned in pairs using Mummer (v4.0) (Marcais et al. 2018) with the parameters -c 500 -b 500 -I 100 -t 6 -maxmatch, then filtered using delta-filter with parameters -1 -i 90 -I 500. The alignments were then used for variation detection with the SyRI pipeline (Goel et al. 2019). Annotation of variations was performed with ANNOVAR (Wang et al. 2010).

### Transcriptome sequencing and identification of specifically expressed genes

Raw RNA-seq data from four tissues (root, stem, leaf, and flower) of *O. vulgare* ‘Hot & Spicy’ and *O. vulgare* were filtered using the Trimmomatic (Bolger et al. 2014) software. Each read’s first ten and last five low-quality bases were removed, and readings shorter than 36 bp were rejected as part of this quality control procedure. Using bowtie v2.0 (Langmead and Salzberg 2012), the resultant clean reads were mapped against coding sequences (CDS) predicted from two haplotype-resolved genomes. Using Trinity (Grabherr et al. 2011), the RSEM v1.3.2 program (https://github.com/deweylab/RSEM) was used to determine fragments per kilobase of exon per million fragments mapped, or FPKM. The DEGs were identified as the ones with log|FC| > 2 and P-value < 0.05.

### Re-sequencing of F_2_ genetic population

Young leaves were gathered from 232 F_2_ individuals as well as from the two parents *O. vulgare* ‘Hot & Spicy’ and *O. vulgare*. The Plant DNA Isolation Kit (DE-0611, http://www.foregene.com) was used to extract DNA. The Truseq Nano DNA HT Sample Preparation Kit (Illumina, USA) was used to create the DNA sequencing libraries, and the Illumina HiSeq2500 platform was used to sequence the samples in accordance with the manufacturer’s instructions. 150 bp paired-end readings were produced from the whole genomes of the parents and the segregating offspring.

### SNP identification

In order to eliminate adapters, read pairs with a proportion of N >10%, and read pairings with low-quality bases (Q ≤ 5) >50%, the sequence reads were processed using Trimmomatic v0.39 (Bolger et al. 2014). The cleaned reads were aligned to two haplotype-resolved genomes hap1 and hap2 using the BWA-MEM2 (v2.2) (Li and Durbin 2009). SNPs and small InDels were mainly detected by GATK (McKenna et al. 2010). According to the localization results of Clean Reads in the reference genome, SAMtools (v1.9) (Danecek et al. 2021) was used to filter redundant reads to ensure the accuracy of the detection results. Then, the HaplotypeCaller (local Haplotypic assembly) algorithm of GATK (v3.8) was used for SNP and InDel mutation detection. Each sample first generated its own gVCF, and then performed the population joint-genotype. We tested the observed segregation pattern of all SNP sites against the expected Mendelian segregation in the F_2_ genetic population using the χ2 test and discarded those sites with distorted segregation (P < 0.05).

### High-density genetic map construction and collinearity analysis

Bin maps and high-density genetic maps were constructed by HighMap which used the developed SNPs bewteen the parents (Li and Durbin 2009). The method of resequencing the entire genome yielded millions of SNP markers of superior quality, surpassing the computational capabilities of existing systems. The three sets of SNP markers were separately fed into Binmarker (v2.3) to construct Bins (https://github.com/lileiting/Binmarkers-v2), which help to minimize the amount of redundant markers. After marker filling and correction, Bin division was performed according to progeny recombination. The samples were arranged neatly according to the physical location of pseudochromosomes hap1 and hap2, and the SNPs between the recombination breakpoints were classified into Bin, and no recombination event was considered to have occurred in Bin. Finally, unfiltered Bin markers were obtained, and Bin was used as the marker for genetic map construction. Bins whose Bin length is less than 5kb and severe segregation (P < 0.00001) were excluded. The collinearity analysis was performed between two high-density genetic maps and two physical maps hap1 and hap2 using a Python script.

### Observation and density of GSTs

The morphology and distribution of the GSTs were evaluated using a stereomicroscope (Leica DVM6, Germany), fluorescence microscope (Leica DM6 B, Germany), and scanning electron microscopy (S-4800, Hitachi, Tokyo, Japan). The leaf area was measured and the GSTs were counted using the ImageJ program. Three plants were used to calculate the density. Finally, the density of GSTs on different tissues in *O. vulgare* ‘Hot & Spicy’ and *O. vulgare*, and GADS, GABS, and GTS traits from F_2_ segregating population.

### Quantitative trait locus mapping analysis

QTL mapping was performed using composite interval mapping (CIM) about the density of GADS, GABS, and GTS with MapQTL v6.0 (Van Ooijen et al. 2009). Significant QTLs were defined as those whose mean values exceeded LOD = 3.0 for the interval mapping mapping approach. A significance level of Signif. < 0.005 was used to the Kruskal-Wallis test. By determining each Bin marker’s physical location on two haplotype-resolved genomes, the QTL intervals in two high-density genetic maps were utilized to identify potential genes. Functional annotations of candidate genes in the QTL intervals were performed using BLASTP against the public databases COG, GO, and KEGG , respectively.

### Candidate gene selection and qRT-PCR analysis

According to the total glandular secretory trichomes density of leaves, we selected 30 individuals of F_2_ genetic population with the highest density and 30 individuals with the lowest density respectively, and constructed 2 mixing pools (HGTS and LGTS). To amplify five candidate genes (*Ovh102G005690*, *Ovh215G004930*, *Ovh215G003840*, *Ovh215G004580*, and *Ovh215G004870*), primers were designed using Primer3 (http://primer3.ut.ee) (Supplementary Data Set 31). Then, qRT-PCR was carried out on two parents and mixing pools cDNAs.

### Statistical analysis

All samples were prepared and analyzed in triplicate, and data were expressed as the mean ± standard deviation. Statistical analyses were performed using the variance (ANOVA) test. Duncan’s test was used to determine the significance of differences between the groups. Differences at P < 0.05 were considered significant. SPSS 18 (SPSS Inc., Chicago, IL, USA) was used for the analysis.

### Accession numbers

The RNA-seq and resequencing data described in this study have been deposited in GenBank of NCBI with BioProjects PRJNA842006 and PRJNA1103877 and in the sequence read archive (SRA) under accession number SRR19383284, SRR19383283, SRR19383282, SRR19383281, SRR19383280, SRR19383264, SRR19383263, SRR19383262, SRR19383260, SRR19383261, SRR19383254, SRR19383253, and SUB14396786.

## Acknowledgments

We thank Xiuping Xu and Ronghua Liang from the Plant Science Facility of the Institute of Botany, Chinese Academy of Science, for their excellent technical assistance in scanning electron microscopy and fluorescence microscopy.

## Author contributions

M.S. and J.M. performed the experiments, analyzed the data and wrote the manuscript.

N.L. helped to evaluate type and density of the parent glandular secretory trichomes.

Y.Z. and J.Z. helped to analyze the density of F_2_ genetic population glandular secretory trichomes. D.W., F.X., and H.B. helped to collect the samples and design the research. L.S. and H.L.was involved in designing the research. M.S., L.S. and H.L. was involved in revising the manuscript. All authors read and approved the manuscript.

## Data availability

Data supporting the findings of this work are available within the paper and its Supplementary Information files. A reporting summary for this article is available as a Supplementary Information file. Genome assembly and annotation have been deposited in GenBank under BioProjects PRJNA842006 and PRJNA1103877.

## Supplementary data

The following materials are available in the online version of this article.

**Supplementary Figure S1.** Construction of F_2_ genetic population between *O. vulgare* ‘Hot & Spicy’ and *O. vulgare*.

**Supplementary Figure S2.** GenomeScope profiles and k-mer distribution analysis of *O. vulgare* ‘Hot & Spicy’.

**Supplementary Figure S3.** The protein-coding genes were predicted in *O. vulgare* ‘Hot & Spicy’ haplotype-resolved genome hap1.

**Supplementary Figure S4.** The protein-coding genes were predicted in *O. vulgare* ‘Hot & Spicy’ haplotype-resolved genome hap2.

**Supplementary Figure S5.** The enrichment analysis of protein-coding genes in *O. vulgare* ‘Hot & Spicy’ haplotype-resolved genome hap1.

**Supplementary Figure S6.** The enrichment analysis of protein-coding genes in *O. vulgare* ‘Hot & Spicy’ haplotype-resolved genome hap2.

**Supplementary Figure S7.** The enrichment analysis of protein-coding genes in *O. vulgare* ‘Hot & Spicy’ haplotype-resolved genome hap1.

**Supplementary Figure S8.** The GC depth and fragment size distribution of Hi-Fi and Hi-C sequencing in *O. vulgare* ‘Hot & Spicy’ haplotype-resolved genome hap2.

**Supplementary Figure S9.** The genome-wide Hi-C heat map of chromosome interactions in two haplotype-resolved genomes hap1 and hap2.

**Supplementary Figure S10.** Circos diagram of two haplotype-resolved genomes hap1 and hap2.

**Supplementary Figure S11.** Cluster analysis of gene families and gene in the haplotype-resolved genomes and other 12 related species.

**Supplementary Figure S12.** Cluster analysis of gene families in two haplotype-resolved genomes and other 12 related species.

**Supplementary Figure S13.** Cluster analysis of specific genes in two haplotype-resolved genomes hap1 and hap2.

**Supplementary Figure S14.** Cluster analysis of contraction genes in two haplotype-resolved genomes hap1 and hap2.

**Supplementary Figure S15.** Cluster analysis of expansion genes in two haplotype-resolved genomes hap1 and hap2.

**Supplementary Figure S16.** Cluster analysis of positive genes in two haplotype-resolved genomes hap1 and hap2.

**Supplementary Figure S17.** KEGG analysis of the intragenic SNP and InDel clustered among two haplotype-resolved genomes.

**Supplementary Figure S18.** KEGG analysis of the structural variation (SV) clustered among two haplotype-resolved genomes.

**Supplementary Figure S19.** Sequencing results of oregano transcriptome sequencing.

**Supplementary Figure S20.** Chromosome coverage depth distribution in two haplotype-resolved genomes hap1 and hap2.

**Supplementary Figure S21.** Variant calling identified all SNPs and InDels between two parents comparing two haplotype-resolved genomes hap1 or hap2.

**Supplementary Figure S22.** Bin markers by illustrated genotype analysis in hap1.

**Supplementary Figure S23.** Bin markers by illustrated genotype analysis in hap2.

**Supplementary Figure S24.** Collinearity comparison between genetic map hap1 and physical map hap1.

**Supplementary Figure S25.** Collinearity comparison between genetic map hap2 and physical map hap2.

**Supplementary Figure S26.** Correlation scatter map between genetic map and physical map.

**Supplementary Method S1.** Haplotype-resolved genomes of *O. vulgare* ‘Hot & Spicy’ sequencing, assembly and annotation.

**Supplementary Data Set 1.** Global statistics of assembly and annotation of two haplotype-resolved genomes.

**Supplementary Data Set 2.** The sequence length of reads in HiFi sequencing of *O. vulgare* ‘Hot & Spicy’.

**Supplementary Data Set 3.** Statistics of the sequencing and assembly results of CSS in *O. vulgare* ‘Hot & Spicy’ genomes hap1 and hap2.

**Supplementary Data Set 4.** Statistical data of various types of Hi-C sequencing in *O. vulgare* ‘Hot & Spicy’ genomes hap1 and hap2.

**Supplementary Data Set 5.** Assembly data statistics of Hi-C sequencing in *O. vulgare* ‘Hot & Spicy’ genome hap1.

**Supplementary Data Set 6.** Assembly data statistics of Hi-C sequencing in *O. vulgare* ‘Hot & Spicy’ genome hap2.

**Supplementary Data Set 7.** Gene prediction result statistics in *O. vulgare* ‘Hot & Spicy’ genomes hap1 and hap2.

**Supplementary Data Set 8.** Statistical table of gene information of *O. vulgare* ‘Hot & Spicy’ genomes hap1, hap2 and related species.

**Supplementary Data Set 9.** Statistics information of gene function annotation in *O. vulgare* ‘Hot & Spicy’ genomes hap1 and hap2.

**Supplementary Data Set 10.** Long terminal repeat (LTR) in *O. vulgare* ‘Hot & Spicy’ genome hap1.

**Supplementary Data Set 11.** Long terminal repeat (LTR) in *O. vulgare* ‘Hot & Spicy’ genome hap2.

**Supplementary Data Set S12.** Microsatellite repeat sequences in *O. vulgare* ‘Hot & Spicy’ genomes hap1 and hap2.

**Supplementary Data Set 13.** Genome information of related species used in comparative genomics analysis.

**Supplementary Data Set 14.** Statistical information of gene family cluster in 13 species.

**Supplementary Data Set 15.** Sequencing data statistics of transcriptome sequencing.

**Supplementary Data Set 16.** Statistics on data mapping of transcriptome sequencing comparison of hap1.

**Supplementary Data Set 17.** Statistics on data mapping of transcriptome sequencing comparison of hap2.

**Supplementary Data Set 18.** Evaluation and statistics of sequencing data of F_2_ genetic population.

**Supplementary Data Set 19.** Comparison of resequencing data between F_2_ genetic population and reference genome hap1.

**Supplementary Data Set 20.** Comparison of resequencing data between F_2_ genetic population and reference genome hap2.

**Supplementary Data Set 21.** SNP statistics of F_2_ genetic population compared with reference genome hap1.

**Supplementary Data Set 22.** SNP statistics of F_2_ genetic population compared with reference genome hap2.

**Supplementary Data Set 23.** InDel statistics of F_2_ genetic population compared with reference genome hap1.

**Supplementary Data Set 24.** InDel statistics of F_2_ genetic population compared with reference genome hap2.

**Supplementary Data Set 25.** Bin screen of F_2_ genetic map hap1.

**Supplementary Data Set 26.** Bin screen of F_2_ genetic map hap2.

**Supplementary Data Set 27.** Information statistics of F_2_ genetic map hap1.

**Supplementary Data Set 28.** Information statistics of F_2_ genetic map hap2.

**Supplementary Data Set 29.** The collinear correlation coefficients between 15 linkage groups and physical map hap1.

**Supplementary Data Set 30.** The collinear correlation coefficients between 15 linkage groups and physical map hap2.

**Supplementary Data Set 31.** Primer sequence of thirty six candidate genes and one reference genes (elf4a).

